# Perceived closeness and autistic traits modulate interpersonal vocal communication

**DOI:** 10.1101/133066

**Authors:** Olivia Spinola, T.A. Sumathi, Nandini Chatterjee Singh, Bhismadev Chakrabarti

## Abstract

Vocal modulation is a critical component of interpersonal communication. It not only serves as a dynamic and flexible tool for self-expression and linguistic information but also plays a key role in social behaviour. Variation in vocal modulation can be driven by individual traits of the individual interlocutors, as well as by factors relating to the dyad, such as the perceived closeness between interlocutors. Accordingly, the current study examines the role of a) individual differences in autism-related traits, and b) perceived closeness between interlocutors on vocal modulation. Since lack of appropriate vocal modulation is often associated with Autism Spectrum Disorders we also focus on autism-related individual traits. The role of these individual and dyad-level factors on vocal modulation is tested for cultural generalizability by conducting this study in three separate samples from India, Italy, and the UK. Articulatory features were extracted from recorded conversations between a total of 85 same-sex pairs of participants and correlated with their self-reported perceived closeness (CR) to the other member of the pair and with the individual Autism Spectrum Quotient (AQ). Results indicated a significant positive correlation between interpersonal closeness and articulation area in all three samples. A significant negative correlation between AQ and articulation area was observed only in the UK sample. This study thus provides novel insights into determinants of interpersonal vocal communication and a test of their cultural generalizability.

## Introduction

Modulation of the human voice is central to interpersonal communication. It serves not only as a dynamic and flexible channel for self-expression and linguistic information but also as a social tool. Consequently humans routinely and volitionally modulate their voice in social contexts like striking up a new friendship or making a compelling argument. A well modulated voice carries considerable information: about the message, speaker, language and even on the emotions of the speaker (Bhaskar, Nandi, & Rao, 2013; Spinelli, Fasolo, Coppola, & Aureli, 2017). Not surprisingly, a number of factors influence context-specific vocal modulation. In acoustic terms, voice modulation is defined as the manipulation of any non-verbal property of the voice including but not limited to F0 and formant frequencies. Recently Pisanski and collaborators (Pisanski et al., 2016/2) distinguished two different types of vocal variations, one being involuntary, and as such automatically elicited by environmental stimuli or endogenous different levels of arousal, and a second, more controlled vocal modulation, that is goal-directed and less dependent on external stimuli, though not necessarily voluntary.

Recently, the context dependency of voice modulation has become a subject of research interest (Pisanski et al., 2016/2). People communicate differently depending on their type of relationship. For instance, the way a person talks to parents is different from the way he or she talks to a friend. Investigating ways to distinguish family from friends through a spontaneous dialogue, Katerenchuk, Brizan and Rosenberg (2014) used OpenSMILE to extract acoustic features from a corpus of native English speech telephone conversations. The OpenSMILE feature extraction process firstly extracts a set of short frame-based low-level descriptors (LLDs) and then applies functionals over these descriptors to extract aggregated features. The feature set included five Low Level Descriptors: 1) Zero Crossing Rate, 2) RMS Energy, 3) F0, 4) Harmonic-to-Noise Ratio, and 5-16) 12 MFCC coefficients.

The authors found a set of spectral features to be discriminative: pitch features indicated an earlier maximum pitch and delta pitch in friends conversations, and results generally suggested that it was possible to distinguish a conversation with friends from one with family on the basis of some low level lexical and acoustic signals. Converging results were reported from another study by Campbell which examined the speech of a single Japanese speaker as she spoke to different conversational partners, finding significant differences in pitch (F0) and normalized amplitude quotient (NAQ)depending on the relationship between the speaker and her conversational partner (Campbell, 2002). These preliminary studies provide evidence and insight on how voice modulation can be affected by context-specific factors.

In addition to context, the role of culture in voice modulation remains scarcely investigated. Cultures may vary in the way they modulate the expression of their feelings. Japanese people for instance seem to exercise considerable control of the display of their own feelings in the face compared to people from western European/American cultures (Ekman et al., 1987; Matsumoto, Takeuchi, Andayani, Kouznetsova, & Krupp, 1998). Similarly there are variations of display rules within culture. People in individualistic cultures tend to see themselves as independent, and authenticity is seen as an ideal goal in communication. They consider consistent behaviour across social situations and interaction partners crucial to maintain the integrity of one’s identity (Noon & Lewis, 1992). In contrast, in collectivistic culture for which the Japanese might be a good representative, appropriate adaptation to one’s interlocutor in the current social context is the ideal, rather than the maintenance of a high degree of consistency across contexts (Safdar et al., 2009). This different emphasis may lead to cultural difference in display rules and might also have an influence in the way vocal modulation occurs across different cultures.

The primary objective of this study was to compare voice modulation in a specific social context across cultures. Past studies of such vocal control or articulation have been directed at investigating the static prosodic characteristics across languages and calculated differences in cues like intonation, rhythm and stress (Mary, 2006; Patel & Daniele, 2003). The term ‘vocal modulation’ by its sheer terminology suggests change - in the context of voice with both time, intensity and frequency. There has, however, been little investigation of the dynamic articulatory features of speech. Given that the speech signal involves the production of sounds at multiple time scales (Poeppel, Idsardi, & van Wassenhove, 2008; Rosen, 1992) which would be represented in voice modulation, we examined different articulatory gestures of speech. Recently, the Speech Modulation Spectrum (SMS) was developed (Singh & Singh, 2008) to study the organisation of articulatory gestures of speech at different time scales. A collection of the different articulatory gestures in a voice signal may be represented as an ‘articulation space’. The ‘articulation space’ provides a spectro-temporal energy distribution of different articulatory gestures and includes both segmental and suprasegmental features. The SMS has been used successfully to compare speech imitation abilities across individuals with and without language talent (Christiner & Reiterer, 2013). Since past research has indicated that voice modulation varies with context we propose the articulation space as an objective measure of social relationship.

To our best knowledge no previous study has used a dimensional approach to test the impact of the social relationship on vocal modulation, or tested the cultural generalisability of such a phenomenon. To take a dimensional approach to evaluate the effect of social relationship, we asked participants to rate how close they felt to another person. Such a ‘closeness rating’ is akin to the widely used metric of ‘social distance’ (O’Connell, Christakou, Haffey, & Chakrabarti, 2013). The literature on social relationship differentiates between subjective closeness (perceptions of relationship quality) and behavioral closeness (degree and variety of interaction) (Aron, Aron, & Smollan, 1992). The ‘closeness rating’ as used in our study relates more to the former than the latter. Considering anecdotal reports and previous related studies, we hypothesized that higher closeness rating for the listener will be associated with an increased number of articulatory gestures by the speaker. Consequently, articulation space and closeness rate were hypothesized to be positively correlated. We expected any such relationship to be true across different cultures, and accordingly tested the generalisability of this relationship in three cultures. Variability in the extent of usage of articulatory gestures in communicative speech can also depend on individual-level factors, and not just on those specific to the dyad and the context. Autism-related traits constitute one such dimension of individual variability. We were interested to examine the impact of autistic traits on the articulation space, given the nature of autism as a disorder commonly associated with deficits in social communication. Anecdotal reports have suggested an ‘autistic monotone’ in describing an absence of context-appropriate vocal modulation in interpersonal communication in individuals with ASD (Bone et al., 2012). Autistic traits exist across a continuum across the population (Baron-Cohen, 2000; Robinson et al., 2011), and thus allows us to examine the impact of individual variability on these traits on the extent of articulatory gestures used in dyadic communication. In line with anecdotal reports, we expected that individuals high in autistic traits would exhibit reduced articulation space, therefore a reduced use of articulatory gestures.

## Methods

### Study Approval

All methods were carried out in accordance with guidelines and regulations included in the official documents of the National Brain Research Centre for the Indian data, University of Reading for the British data and of the Universita’ degli studi di Milano Bicocca for the Italian data. All experimental protocols were approved by the Research Committee of the University of Reading, Universita’ degli studi di Milano Bicocca and National Brain Research Centre.

### Participants

170 healthy volunteers from three countries (43 pairs of participants from UK, 22 pairs from India and 20 pairs from Italy) participated in this study (See Table 2).

**Table 1.** Correlations

| Site | N | Age<br>(mean,SD) |
| --- | --- | --- |
| Reading, United Kingdom | 86 (36 males, 50 females) | 20.1; 2.4 |
| Gurgaon, India | 44 (18 males, 26 females) | 24.6; 3.66 |
| Rome, Italy | 40 (20 males, 20 females) | 21.2; 2.2 |

**Table 2.**
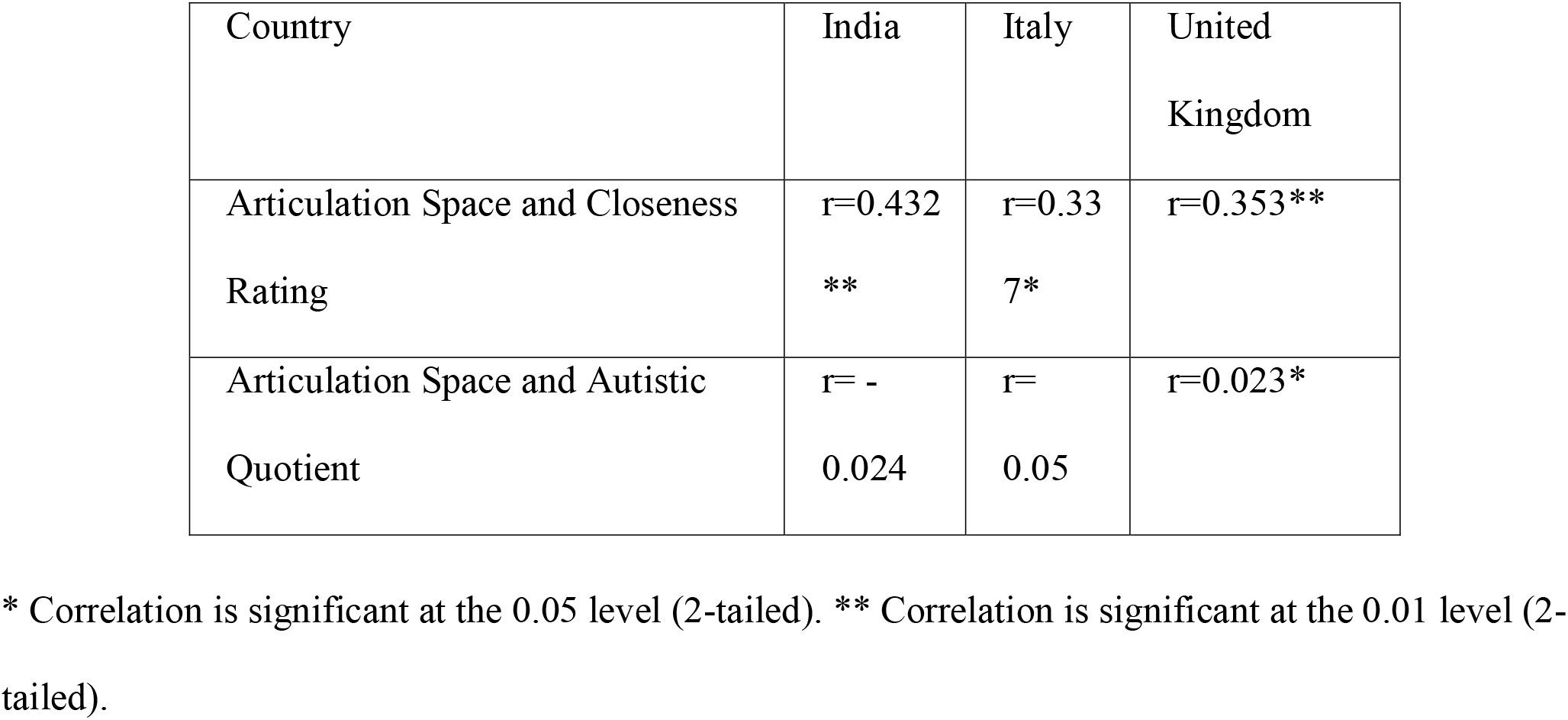
Participants Details for each sample

### Procedure

Participants were asked to come along with another same-gender person, which could be either a friend or an acquaintance. Each participant was asked to rate his/her perceived closeness to the other member of the pair through a Closeness Rating scale on a 10-point Likert scale (where closeness rating of 0 indicated very low closeness and 10 high closeness). Participants were not allowed to see each other’s Closeness Rating. All participants were also asked to fill in the Autism Spectrum Quotient (Baron-Cohen, 2000). They were required to sit in front of each other in a silent room with a voice recording device in the middle. Once participants were comfortably seated, each participant was given one of two coloured images (abstract paintings, attached) printed on paper and was asked to describe the image to the other participant in as much detail as possible for around 2:30 minutes each. The experimenter then left the room. Each pair was instructed to speak one by one, to avoid overlapping voices and making background noises. The participant’s speech was recorded by the ZOOM H1 Handy recorder in India, and an iPhone in UK and in Italy. The distance of the recorder was kept constant for all recordings. Italy and UK participants were asked to speak in their native language while the Indian participants were asked to speak in English.

### Analysis

All participants’ speech was manually listened to carefully and cleaned using Goldwave (version 5.69) and resampled as PCM signed 16 bit mono, 22050 Hz sampling rate in WAV format. The speech data was edited manually and non-speech utterances such as laughs, cough etc. were removed. The amplitude of the waveforms was normalized to −18db for all speech data. Noisy speech data was excluded from the analysis. Speech Modulation Spectra (Christiner & Reiterer, 2013; L. Singh & Singh, 2008) were calculated for each participant using custom developed code developed in MATLAB R2011a. Statistical analysis was carried out using SPSS(version 14). Finally articulation space, closeness rating and AQ measures were used for further analysis in all three datasets. The length of the cleaned speech recordings varied in India 40.6s to 185.1s(mean=119.2, SD=27.1), Italy 95.5s to 179.7s(mean=130.2, SD=18.5) and UK 34.8s to 232.5s (mean=112.0,SD=25.0). 35s to 232s (mean=119.2s s.d.=25.2s. Articulatory features were calculated for each individual using the SMS (Speech Modulation Spectrum)method implemented in custom-written MATLAB code (L. Singh & Singh, 2008; N. C. Singh & Theunissen, 2003). (L. Singh & Singh, 2008) developed this novel spectral analysis technique, called Speech Modulation Spectrum to study the organization of articulatory gestures as a metric of speech motor skills.

The first step of this analysis involves using speech samples from each participant to calculate a spectrogram. The spectrogram is a time-frequency representation of the speech signal and offers a visual display of fluctuations in frequency and time (see Figure 1), described respectively as spectral and temporal modulations.

**Figure 1.**
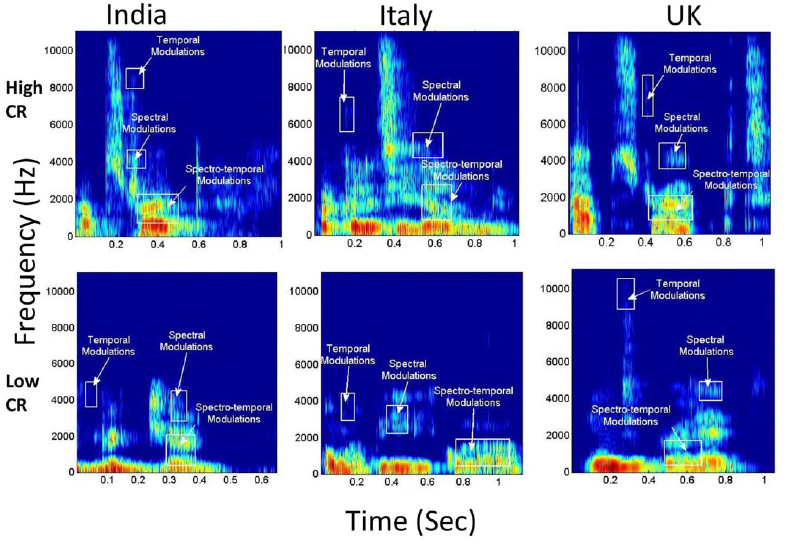
Representative spectrogram of speech samples of High and Low closeness rate(CR) for all three groups

As shown in Figure 1, spectral modulations (wf) are energy fluctuations across a frequency spectrum at particular times, whereas temporal modulations (wt) are energy fluctuations at a particular frequency over time. Based on the rate of fluctuation, spectro-temporal modulations have been proposed to encodes both suprasegmental and segmental features. A 2-D Fourier transform of the spectrogram yields a probability distribution of these different articulatory features and is called the Speech Modulation Spectrum (N. C. Singh & Theunissen, 2003). In a typical speech modulation spectrum, the central region between 2-10 Hz carries supra-segmental information and encodes syllabic rhythm. The side lobes between 10-100 Hz carry information about segmental features. Formant transitions are encoded between 10-40 Hz, and place of articulation information is found between 40-100 Hz ((Blumstein & Stevens, 1980; Tallal, Stark, & Mellits, 1985). As the modulation spectrum goes from 1-100 Hz, the amplitude fluctuations of a sound become faster and go from syllabic to vowel-like to plosive-like segments ((N. C. Singh & Theunissen, 2003)). The speech modulation spectrum thus plots an “articulation space,” which depicts how energy or ‘power’ is distributed in different articulatory features of spoken language, namely syllabic rhythm, formant transitions, and place of articulation (see Figure 2).

**Figure 2.**
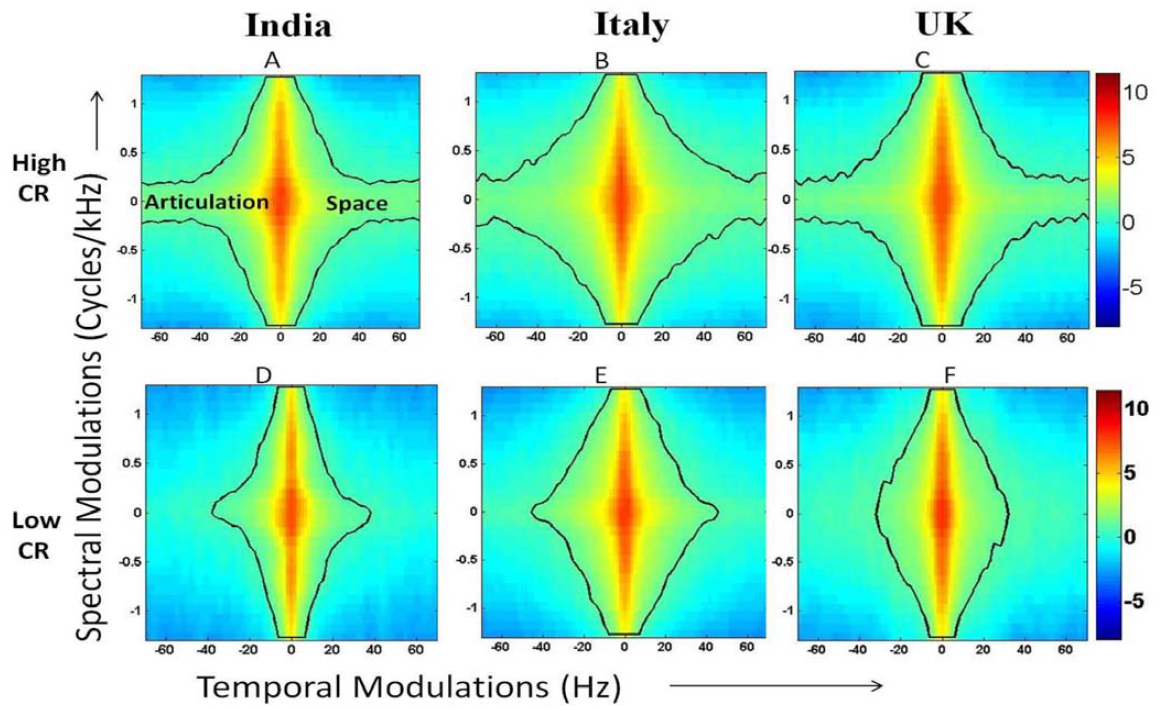
Representative Speech modulation spectrum of High and Low closeness rate(CR) for all three groups.

The articulation space is estimated by calculating the total contour area of the spectro-temporal modulations that encompass 99.9% of the total energy. The speech modulation spectrum was plotted for each participant and the contour area (hereafter referred to as ‘articulation space’) was estimated by counting the total number of pixels within the contour which covers 99% energy of the speech (Christiner & Reiterer, 2013; L. Singh & Singh, 2008) from 1 to 100 Hz. We investigated the entire contour area from 1 to 100Hz which corresponds to all the articulatory features mentioned above (detailed description in (L. Singh & Singh, 2008). Previous studies used this method and demonstrated its construct validity by testing its correlation with other behavioral measures, such as speech motor functions (Christiner & Reiterer, 2013; Sullivan, Sharda, Greenson, Dawson, & Singh, 2013). In the current study, the articulation space was correlated with closeness ratings and AQ.

#### Outlier check

Cook’s distance was calculated to determine the outlier in articulation space and length of speech recordings.

#### Correlations

Bivariate correlations were computed between articulation space and length of speech recordings to test whether the duration of the speech affects the articulation space or not. Separate bivariate correlations were computed with articulation space and interpersonal closeness and autistic traits to test the two key study hypotheses. All bivariate correlations partialled out the effect of gender and the duration (length of speech).

## Results

### Energy Distribution and Articulation Space Area

The spectrogram of vocalizations from the participant’s speech sample represents the spectral, temporal and the spectro-temporal modulations which are highlighted in Figure 1.

There we found difference in the energy distribution in the spectro-temporal modulations between the high and low closeness rate participants. Speech Modulation Spectrum (SMS) images(See Figure 2) were plotted for all participants. The speech modulation spectrum was used to demonstrate the differences in the articulatory features between the high and low closeness rate participants across all three groups. The color bar indicates the energy distribution of articulatory features. A difference in the articulation space area (0–100Hz)was found between the high and low closeness ratings people across all three groups. So this measure was used to correlate with other measures namely closeness rate and autistic quotient (AQ).

### Closeness Rating and Articulation Space Area

The bivariate correlation (See Figure 3) showed that there was a moderate, positive correlation between closeness rate (CR) and articulation space area which was statistically significant in all three cultures.

**Figure 3.**
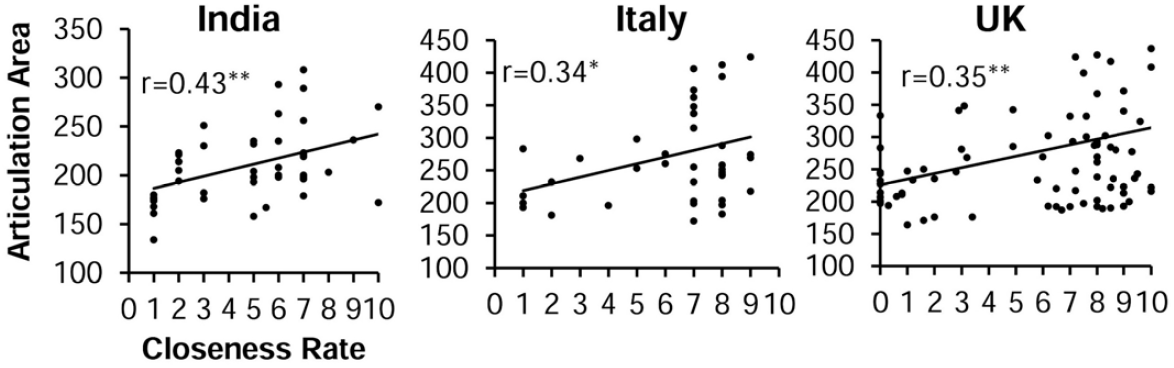
Correlation between Closeness rate and articulation area.

This indicates that the articulation space significantly increases when the closeness rate increases in all three countries (India r=0.43, p<0.001; Italy r=0.34, p<0.01; UK r=0.35, p<0.001)(See Table 1).

### Autistic Quotient (AQ) and Articulation Space area

The bivariate correlation between the articulation space area and autistic quotient (AQ) showed that there was a negative correlation (See Figure 4) which was statistically significant only in the UK group (r=-0.23, p<0.01).

**Figure 4.**
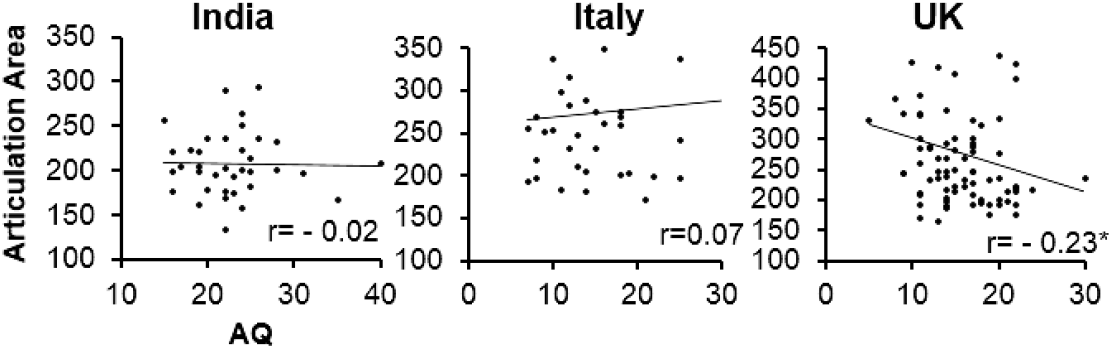
Correlations between Autistic Quotient and Articulation Area

The same trend was seen in the India group but was found to be not significant. The Italy group did not show any trend (See Table 1).

### Data Exclusion

3 pairs from the India dataset and 1 pair from the UK dataset were excluded from this analysis si ce their AQ measure was not available.

## Discussion

This study investigated the variability in articulation space during dyadic vocal communication in relation to two factors: the self-reported closeness of the interactants, as well as individual autistic traits. These relations were tested for cultural generalisability by running the same study in three different cultures. The key finding was a positive association between the closeness rating and articulation space: individuals used greater articulatory gestures when they spoke to those who they rated high on closeness ratings, compared to others who they rated low. This finding was consistent across all three cultures studied. At an individual level, autistic traits were found to be inversely related to the articulation space. Interestingly, this result appears to be culture-specific, as it was observed only in the UK sample.

Anecdotal accounts suggest that people use greater modulation of their voice during speaking to familiar others compared to strangers (Katerenchuk et al., 2014) This aspect of deliberate vocal control has been noted in other nonhuman primates and predates human speech (Pisanski et al., 2016/2). From a functional perspective, articulation space can arguably contain informative cues about group membership, and hence might subserve social bonding processes. Communication accommodation theory suggests that one of the affective motives of interpersonal vocal communication is to manage and regulate social distance (Dragojevic, Gasiorek, & Giles, 2016). Increased closeness was associated with a greater articulation area, which suggests that a) more information is being communicated with a closer other, through incorporation of greater non-verbal signals, or/and b) individuals are more inhibited by social norms when talking with someone who they are not close to, and thus reduce the number/extent of their articulatory gestures. The current dataset does not allow us to discriminate effectively between these two possibilities.

This relationship between closeness and articulation space was found to be similarly robust in all three cultural settings. This observation, despite the variability of languages used in the different samples (English in UK and India, Italian in Italy), suggests that this relationship is relatively independent of language and cultural environment. This finding extends the evidence base for cross-cultural generalisability that has been noted for several other social cognitive processes, such as recognition of facial and vocal expressions of certain emotions (Elfenbein & Ambady, 2002; Scherer, Banse, & Wallbott, 2001). Beyond factors specific to the dyad, such as how close an individual felt toward another, the impact of individual variation was measured, in line with an approach suggested by ((O’Connell et al., 2013). Individual variation in autism-related traits were found to be negatively related to the area of the articulation space only in the UK sample and not in the other two samples. Autism has been associated with atypical production of and wider variation in prosody (Bone et al., 2012; Joshua John Diehl & Paul, 2011; Joshua J. Diehl, Watson, Bennetto, McDonough, & Gunlogson, 2009; Filipe, Frota, Castro, & Vicente, 2014; Peppé & McCann, 2003). While this result supports reports of atypical intonation and anecdotally ‘monotonic’ voice in individuals with ASD, an increased pitch range has been noted in ASD in paradigms using single word utterances as well as narratives (Joshua John Diehl & Paul, 2011; Joshua J. Diehl et al., 2009; Filipe et al., 2014; Sharda et al., 2010). It is worth noting that articulation space captures a wider set of acoustic features than pitch range per se. The observed cultural difference in this association between articulation area and autistic traits might arise due to differences in baseline levels of vocal modulation in different cultures. Such differential ‘baseline’ levels of vocal modulation in different cultures might be related to differences in the structure of the languages, e.g. in English it is possible to ask a question using the sequence of words, without the need for an intonation, such as ‘Have you eaten?’. In contrast, in Italian, the interrogative nature of the sentence ‘Hai mangiato?’ is communicated by tonal difference alone: uttering this sentence in a flat tone will be equivalent to making a factual assertion of ‘you have eaten’. However, in view of the fact that the Indian participants also spoke in English, the observed differences can not be attributed solely to the differences in the languages per se.

Several sources of individual and dyadic variation remain unmeasured in this current study, and need to be investigated in future studies. Gender is one such variable, which accounts for significant differences in vocal modulation in the interpersonal context. The current study focused only on same-gender pairs, and thus did not explore this dimension of variability.

In sum, this study found the impact of closeness on vocal modulation in interpersonal communication, demonstrating that a greater closeness was associated with more modulation, across different cultural and language settings. This study also found that autism-related traits were associated negatively with the extent of such vocal modulation, but only in one of the three cultural settings. Future studies should extend these paradigms to include individuals with clinical deficits in social communicative abilities, such as those with ASD, to test the potential culture-specificity of certain phenotypic features of ASD.

## Acknowledgements

The authors wish to thank Lauren Sayers, Eleanor Royle, Bethan Roderick for helping with collection of data in the UK, Emanuele Preti for helping with organizing and collecting data in Italy, Angarika Deb and NBRC students for helping with the data collection in India. BC acknowledges support from the Leverhulme Trust and the Medical Research Council UK.

## Author contributions statement

BC, OS, NCS designed the study; OS, TAS, BC conducted the experiments; TAS, NCS analysed the results. All authors co-wrote and reviewed the manuscript.

## Additional information

**Competing financial interests**

The authors declare no competing financial interests.

